# Ventral pallidal GABAergic neurons drive consumption in male, but not female rats

**DOI:** 10.1101/2024.04.30.591876

**Authors:** Alexandra Scott, Anika Paulson, Collin Prill, Klaiten Kermoade, Bailey Newell, Jocelyn M. Richard

## Abstract

Food intake is controlled by multiple converging signals: hormonal signals that provide information about energy homeostasis, but also hedonic and motivational aspects of food and food cues that can drive non-homeostatic or “hedonic “feeding. The ventral pallidum (VP) is a brain region implicated in the hedonic and motivational impact of food and foods cues, as well as consumption of rewards. Disinhibition of VP neurons has been shown to generate intense hyperphagia, or overconsumption. While VP gamma-Aminobutyric acidergic (GABA) neurons have been implicated in cue-elicited reward seeking and motivation, the role of these neurons in the hyperphagia resulting from VP activation remains unclear. Here, we used Designer Receptors Exclusively Activated by Designer Drugs (DREADDs) to activate or inhibit VP GABA neurons in sated male and female rats during chow and sucrose consumption. We found that activation of VP GABA neurons increases consumption of chow and sucrose in male rats, but not female rats. We also found that, while inhibition of VP GABA neurons tended to decrease sucrose consumption, this effect was not statistically significant. Together, these findings suggest that activation of VP GABA neurons can stimulate consumption of routine or highly palatable rewards selectively in male rats.

## INTRODUCTION

Food intake is controlled by multiple converging signals: the primary attributes of food (i.e. taste and smell), environmental cues that attract individuals to the food, and hormonal signals that provide information about energy homeostasis. Neural circuits integrate these cues and signals to initiate food-seeking and consumption (Morton et al., 2014). Excessive cue reactivity can lead to eating beyond the caloric need for energy homeostasis. The ventral pallidum (VP) is implicated in controlling both food consumption and cue-elicited reward seeking. Disinhibition of the VP increases food intake in rats (Stratford and Kelley, 1999; Stratford and Wirtshafter, 2013), especially of food high in fat (Covelo et al., 2014). The VP is also an essential node in the neural circuitry that underlies motivation to food/reward-seek. VP neurons are known to respond to cues predictive of reward and these neuronal responses encode the motivational value of said cues (Ahrens et al., 2016; Richard et al., 2018, 2016; Smith et al., 2011; Tindell et al., 2005). Recent research has begun to probe which VP neuronal cell-types contribute to consumption and reward seeking. The VP contains 3 major neuronal subtypes: cholinergic, glutamatergic, and gamma-Aminobutyric acidergic (GABA). VP glutamatergic neurons are excited by aversive stimuli and constrain reward seeking (Faget et al., 2018; Tooley et al., 2018). In contrast, many VP GABA neurons are excited by rewards and Pavlovian reward-paired cues (Stephenson-Jones et al., 2019). Recently we found that VP GABA neurons encode the incentive motivational value of cues, and causally contribute to cue-elicited reward-seeking (Scott et al., 2023), yet it remains unclear whether these same neurons are responsible for the hyperphagia produced by hyperactivation of the VP.

Recent work has found mixed effects of VP GABA manipulations on food-seeking and consumption. Chemogenetic inhibition of VP GABA neurons decreases food-seeking (i.e. operant responding) of chow and palatable food (banana pellets) in food-restricted rats (Farrell et al., 2021), whereas inhibition of VP GABA neurons in sated rats reduces palatable food-seeking but not chow-seeking behavior. In contrast, inhibition of VP GABA neurons fails to significantly reduce non-operant consumption of palatable food (Farrell et al., 2021). Prior work also found that activation of VP GABA neurons does not increase consumption of chow in food-restricted mice (Y.-D. Li et al., 2020). Together, these findings might suggest that VP GABA neurons control food-seeking but not consumption, which would be surprising considering the prior work discussed above (Covelo et al., 2014; Cromwell and Berridge, 1993; Stratford and Wirtshafter, 2013). Yet, the internal state of animals is known to impact VP GABA responses to reward and reward-seeking (Farrell et al., 2022, 2021; Stephenson-Jones et al., 2020). Additionally, VP manipulations have been shown to have differential influence over different macronutrients (Covelo et al., 2014). Overall, it is still unclear how manipulation of VP GABA neurons in sated animals affects consumption of home chow or rewards of single macronutrient origin. Furthermore, the prior work has not disaggregated testing subjects by sex.

Here we used chemogenetic methods to test whether VP GABA neuron activation or inhibition affects consumption of chow or 10% liquid sucrose in sated male and female rats. We found that VP GABA neuron activation increases both chow and sucrose intake in a sex-specific manner. In contrast, we found that inhibition of VP GABA neurons does not significantly change either home chow or sucrose intake in sated rats, however rats did tend to consume less sucrose when VP GABA neurons were inhibited. These results suggest that while overactivation of VP GABA neurons can drive increased consumption in sated male rats, these neurons may be less important for basal consumption.

## MATERIALS AND METHODS

### Subjects

Male and female Long Evans rats (n = 62, 32 male, 30 female; Envigo), weighing 250–275 grams at arrival, were individually housed in a temperature- and humidity-controlled colony room on a 14/10 hr light/dark cycle. All experimental procedures were approved by the Institutional Animal Care and Use Committee at the University of Minnesota and were carried out in accordance with the guidelines on animal care and use of the National Institutes of Health of the United States.

### Surgeries

During surgery, rats were anesthetized with isoflurane (5%) and placed in a stereotaxic apparatus, after which surgical anesthesia was maintained with isoflurane (0.5–2.0%). Rats received preoperative injections of carprofen (5 mg/kg) for analgesia and cefazolin (70 mg/kg) to prevent infection. Syringes for viral delivery were aimed at the ventral pallidum (VP) using the following coordinates in comparison to bregma: +0.3 mm AP, +/− 2.3 mm ML, −8.3 mm DV. To achieve cell-type specific expression of designer receptors exclusively by designer drugs (DREADDs) 0.60-0.80 uL of a 1:1 mixture of AAV8-GAD1-cre (8.29×10^13; University of Minnesota Viral Vector Core) and cre-dependent DREADD virus was injected bilaterally into VP. Gq DREADD virus (n=23, 12 males, 11 females: AAV8-hSyn-DIO-hM3Dq-mCherry, ≥ 4×10¹² vg/mL, Addgene; Watertown, MA, USA) was mixed with GAD1-cre to express excitatory designer receptors in VP GABA neurons. Gi DREADD virus (n=25,13 males, 12 females: AAV8-hSyn-DIO-hM4Di-mCherry, ≥ 1×10¹³ vg/mL, Addgene; Watertown, MA, USA) was mixed with GAD1-cre to express inhibitory designer receptors in VP GABA neurons. For external controls (n=14, 7 males, 7 females) 0.6-0.8 uL of a 1:1 mixture of GAD1-cre and mCherry virus (AAV8-hSyn-DIO-mCherry, ≥ 1×10¹³ vg/mL; Addgene, Watertown, MA, USA) was injected bilaterally into VP. Virus was delivered to VP through 33-gauge injectors at a rate of 0.1 μl per min. Injectors were left in place for 10 min following the infusion to allow virus to diffuse away from the infusion site. Rats recovered for at least one week before beginning handling. Rats recovered for at least 4 weeks before training to allow time for sufficient viral expression.

### DREADD Ligand and Administration

Injections of DREADD ligand were administered on test days through intraperitoneal (IP) injection. DREADD ligand, JHU37160 (J60: HB6261, Hello Bio, Princeton, NJ, USA; Bonaventura et al., 2019) was dissolved in saline and diluted to a concentration of 0.05 mg/ml and 0.50 mg/ml. Ligands were injected at 1 ml/kg of volume to body weight.

### Behavioral Testing

#### Chow Consumption Testing

Rats were habituated to the testing environment 3 days before DREADD ligand or saline injection test days. The testing environment consisted of individually housed cages with corn cob pellet bedding and water. On test days, chow (Teklad Global 18% Protein Rodent Diet, pellets, 2818, Envigo) was pre-measured at 20 grams for each subject. On separate test days, counterbalanced for order, rats received IP injections of 0, 0.05, or 0.5 mg/kg J60, and were immediately placed in test cages. One group (n=28) had the chow placed into the cage at the time of injection, meaning these animals had an hour and a half for consumption. Another group (n=34) received chow 30 minutes post-injection, meaning that these animals had an hour for consumption. The final amount of chow was recorded, and the amount consumed was calculated. Observation days followed each test day to ensure that DREADD ligand did not have any lingering consequences on behavior.

#### Sucrose Consumption Testing

10% sucrose was initially provided to subjects overnight prior to the start of the experiment to reduce neophobia. Rats were then habituated to the testing environment 3 days before DREADD ligand or saline injection test days. The testing environment consisted of individually housed cages with corn cob pellet bedding and a two-bottle top with water and sucrose bottles. Bottle placement was balanced across subjects. On test days, water and sucrose bottles were weighed before the test. After IP injection was given, rats were immediately placed in the testing chambers. One group (n=15) received water and sucrose availability at the time of injection, meaning that the animals had access to water and sucrose for an hour and a half. Another group (n=22) received access to water and sucrose 30 minutes post-injection, meaning the animals had access to water and sucrose for an hour. The amount of water and sucrose was recorded after 90 minutes, and the amount consumed was calculated. Observation days followed each test day to ensure that the DREADD ligand did not have any lasting impacts on behavior. Injection order was counterbalanced across animals. Sucrose preference was calculated as the difference between the amount of sucrose consumed and the amount of water consumed, divided by the total amount of liquid consumed.

### Histology

#### Region specific DREADD expression validation

To confirm that DREADD virus was expressed in VP, 40um sections were imaged on a wide field fluorescent microscope (Olympus MV×10). We verified the location of DREADD expression in all animals (n=62) based on anatomical markers of VP (anterior commissure, lateral ventricle size). 7 animals were excluded for lack of bilateral DREADD expression. To quantify the regional selectivity of DREADD virus expression we further examined tissue from the VP as well as the regions anterior to the VP (+0.72 to 2.28 mm ahead of bregma) and posterior to the VP (−0.72 to −1.72 mm mm behind bregma) of representative rats (n=8; n=4, 2 males, 2 females of 600 nl viral injections; n=4, 2 males, 2 females of 800 nl viral injections). All images were acquired with the same acquisition settings. The QUINT workflow (EBRAINS) was then used to assess the 3D spread of virus expression. Briefly, the 2D images of each section from each animal were matched to atlas (Waxholm Space Atlas v4) segmentations in the QuickNII software. Then, masks of mCherry expression, of each section/animal, were made in the iLatsik software, with labels differentiating between background and expression. Finally, quantification of regional expression was calculated with atlas segmentations and masks of sections in the Nutil software. The final output contained all atlas brain regions in our sections and the amount of mCherry expression (pixel quantification) in each region.

#### DREADD Functional Validation

Rats from each DREADD group were administered either 0, 0.05, or 0.50mg/kg J60 in saline solution 1 hour prior to perfusion. Rats were deeply anesthetized with pentobarbital and perfused intracardially with 0.9% saline followed by 4% paraformaldehyde. Brains were removed, post-fixed in 4% paraformaldehyde for 4–24 hr, cryoprotected in 20% sucrose for >48 hr, and sectioned at 40 um on a microtome for cFos immunofluorescence. Briefly, sections were washed in PBS, blocked in Normal Donkey Serum, and incubated in the cFos primary antibody (1:2500 rabbit anti-cFos, Cell Signaling Technology, 2250s) overnight at room temperature. Sections were then washed in PBS and incubated in the fluorescent conjugated secondary antibody (1:250 Alexa 488 Donkey anti-rabbit, Invitrogen # A-21206) for 2 hours at room temperature. Sections were washed in PBS, wet-mounted on coated glass slides in PBS, air-dried, and coverslipped with Vectashield mounting medium with DAPI. Imaging of the tissue was done on a confocal microscope (Nikon A1R, University Imaging Centers at UMN) at 20x with 2x digital zoom. The exact same acquisition settings were used on all tissue samples. HALO software (Indica Labs) was used to quantify the amount of DREADD expressing cells (mCherry positive cells) that colocalized with the early activation gene, cFos. Briefly, total cell number and location within an image was determined by nuclear staining (DAPI). mCherry and cFos were assigned their respective fluorophores within the software and the software compared the location of each fluorophore to the location of the DAPI labeled cells to confirm fluorophore expression is associated with a self-defined nuclei and cell size. HALO analysis software and software settings were optimized for the tissue from all animals and then settings were used to batch analyze cell counts of mCherry and cFos positive cells.

### Statistical Analysis

Statistical significance was measured by fitting a linear mixed model to examine the effects of the designer drugs within DREADD groups (i.e., Gq, Gi, or mCherry), with fixed effects for sex and ligand dose and a random effect of rat. When significant main effects or interactions were found pairwise comparisons were run with Sidak corrections for multiple comparisons. To examine the specificity of the effects of ligand in male Gq rats we fit linear mixed models with fixed effects for virus and ligand dose, and a random effect of rat on Gq and control males. Quantification of DREADD expression spread was also assessed by fitting a linear mixed model to examine the effects of injection volume on viral spread and expression of DREADD virus. Linear mixed models were fit with fixed effects for sex, volume and brain region, and a random effect of rat.

## RESULTS

Here we assessed the impact of chemogenetic activation or inhibition of VP GABA neurons on sucrose and chow consumption in *ad libitum* fed male and female rats. In addition to verifying that all included rats had bilateral DREADD virus expression centered in VP, we conducted more detailed analysis of 8 randomly selected Gi and Gq brains to determine to extent of viral expression spread to adjacent brain regions (Figure 1A-B). Most fluorescence (81.2 +/− 4.86% of objects, a proxy for cell count) was restricted to the VP. We then identified additional atlas regions outside VP in which more than one rat had greater than 5% of their fluorescent object counts. We observed expression exceeding this benchmark in 3/8 rats in the corticofugal tract and corona radiata (8.72 +/− 4.41% mean +/− SEM), and in 4/8 rats in the caudate (8.60 +/− 2.52%). The next region with the most extra-VP expression was the bed nucleus of the stria terminalis (BNST; 1.51 +/− 0.67%), in which no rats had more than 5% of their fluorescent objects. Overall, we did not observe substantial expression in all rats in any specific brain region other than the VP. These results suggest that our behavioral findings are the result of chemogenetic manipulation of VP GABA neuron activity, and not due to changes in activity in adjacent brain structures. We also assessed the functional impact of the DREADD manipulations by assessing cFos expression (Figure 1C-D). In rats expressing Gq DREADD we found that both doses of J60 significantly increased the proportion of mCherry cells expressing cFos (Figure 1C-D; main effect of ligand, F(2,6) = 11.45, p = 0.0089). Importantly for the interpretation of the behavioral results described below, sex had no effect on cFos expression (main effect of sex, F(1,6) = 0.41, p = 0.55) and we observed no interaction between sex and the effect of the ligand (sex by ligand interaction, F(2,6) = 0.12, p = 0.89). We observed no effect of ligand on cFos expression in rats expressing Gi DREADD (main effect of ligand, F(2,6) = 0.18, p = 0.83), and no effect of sex (F(1,6) = 0.02, p = 0.88) or interaction of sex and ligand (F(2,6) = 0.52, p = 0.62). This is likely due to low basal levels of cFos under control conditions.

**Figure 1.**
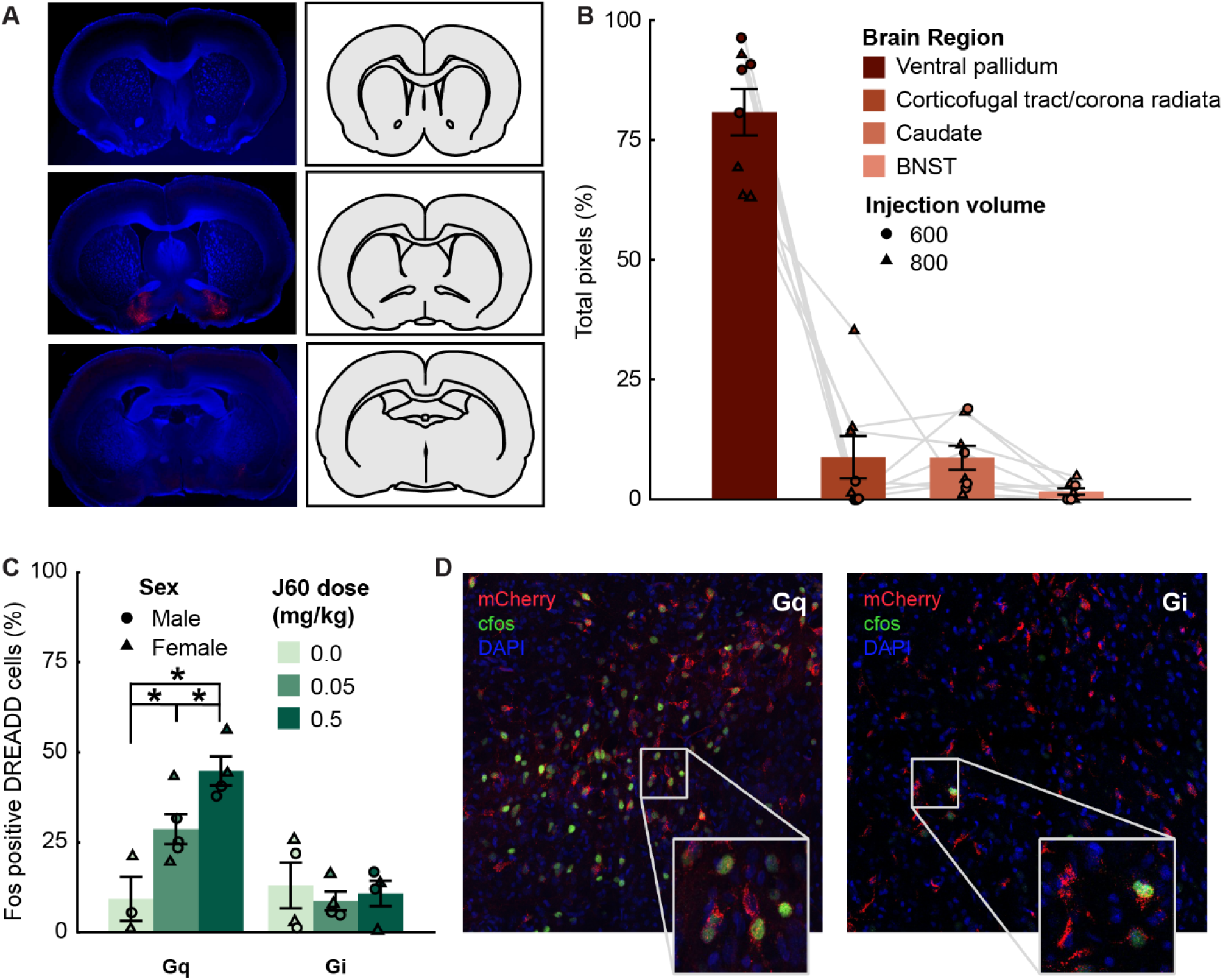
Characterization of DREADD localization and functional impact. (A) Representative section from striatum (top), ventral pallidum (middle), and septum (bottom), used for analysis in QUINT workflow. (B) Output results of QUINT workflow (n=8), showing all brain regions where mCherry (DREADD virus reporter) was detected in rats that received 600 nl (n=4, circles) and 800 nl (n=4, triangles) injections. The VP atlas region was where 81.2 +/− 4.86% of objects (proxy for cell count) were detected by analysis software. LME showed a significant effect of brain region (F(3,28)=113.34, p < 0.001), on spread of virus, with follow-up comparisons indicating that significantly more expression was found in VP was significantly greater in expression compared to other brain regions that were found to have expression in multiple animals (corticofugal tract and corona radiata 8.72 +/− 4.41%, caudate putamen 8.60 +/− 2.52%, bed nucleus of the stria terminalis 1.51 +/− 0.67%). (C) Percent of DREADD-positive cells expressing c-Fos after 0, 0.05 or 0.5 mg/kg J60 in male (circles) and female (triangles) rats expressing Gq or Gi DREADD. Ligand injections increased the % of DREADD-expressing cells that expressed cFos in the Gq group but not the Gi group. * p < 0.05 (left). D) Representative VP images from a Gq male (left) and a Gi male (right) injected with 0.5 mg/kg J60.

### Activation of VP GABA neurons affects consumption of chow and sucrose in a sex-biased manner

When we examined the impact of Gq DREADD chemogenetic activation of VP GABA neurons on chow and sucrose consumption, we found effects that varied by sex (Figure 2). In Gq rats tested with chow (n=17, 8 females, 9 males; Figure 2C) we found a significant effect of ligand dose (F(2, 19.49) = 5.09, p = .01), sex (F(1, 26.19) = 13.68, p = .002) and a significant sex by ligand interaction (F(2, 23.42) = 6.12, p = .006). Pairwise comparisons indicated that the high dose of J60 increased chow consumption in males relative to the saline condition and the low dose condition. In contrast, J60 DREADD ligand had no effect on consumption in female Gq rats. These results indicate that activation of VP GABA neurons selectively impacts chow consumption in male rats.

**Figure 2:**
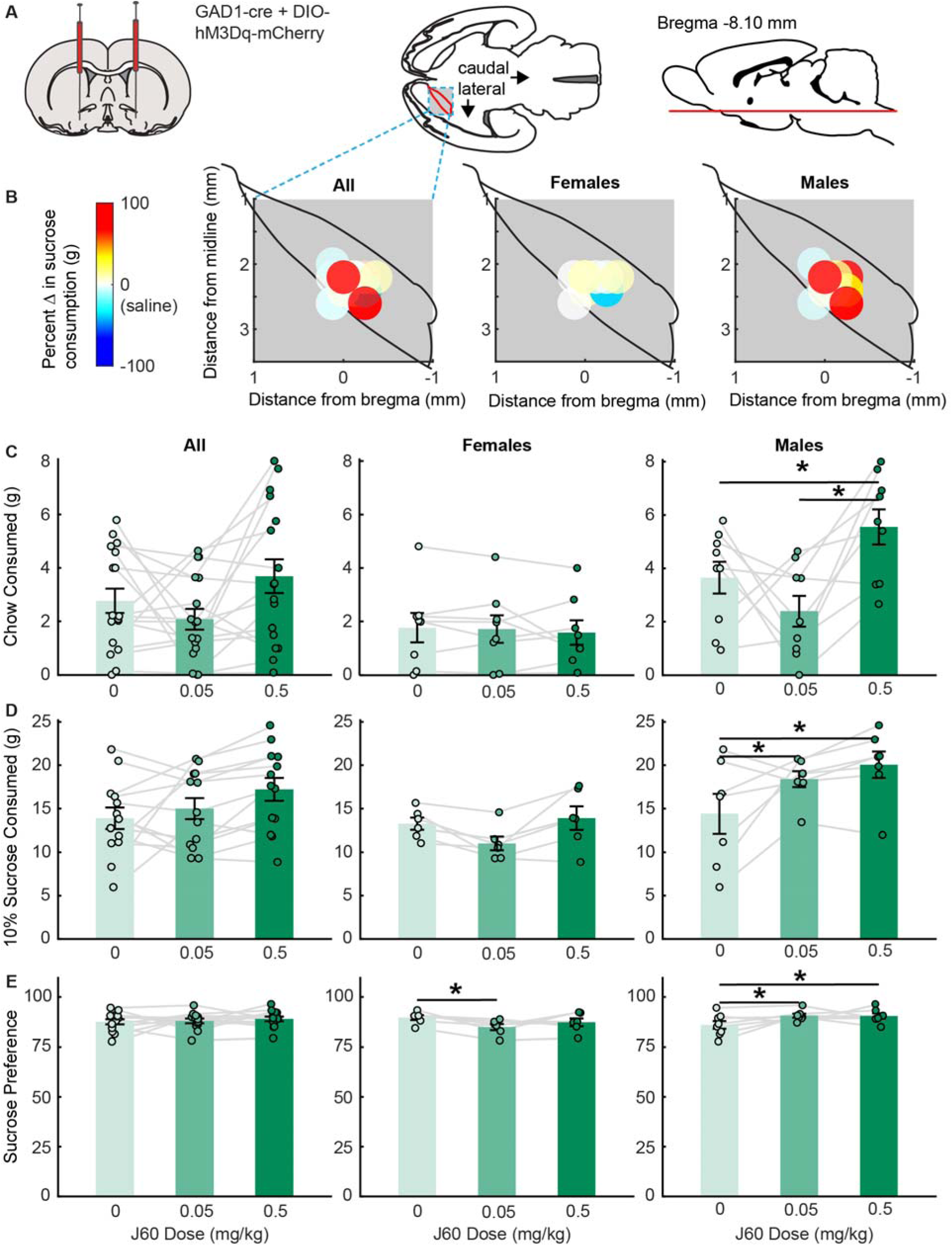
VP GABA activation increases consumption of chow and sucrose in male rats. (A) A mixture of GAD1-cre and DIO-hM2Dq-mCherry was injected bilaterally into the VP. (B) Colormaps show viral placement (brightest point of expression) a horizontal cross section of the VP and behavioral changes from baseline following a high dose (0.5 mg/kg) of J60, for all Gq rats (left), female Gq rats (middle), and male Gq rats (right). Chow consumption (C), sucrose consumption (D), and sucrose preference (E) following injections of saline (light green), low dose J60 (medium green) and high dose J60 (dark green) for all Gq rats (left), female Gq rats (middle), and male Gq rats (right). Bar plots and error bars indicate mean +/− SEM. Dots and lines represent individual rats. Asterix indicate p-values less than 0.05 for Sidak corrected pairwise comparisons.

When we examined the impact of chemogenetic activation of VP GABA neurons on sucrose consumption (n=13, 6 females, 7 males; Figure 2B and D) we observed similar sex-dependent effects. We found significant effects of ligand dose (*F*(2, 22) = 4.27, *p* = .03) and sex (*F*(1, 11) = 9.60, *p* = .01), and a ligand by sex interaction (*F*(2, 22) = 4.40, *p* = .03). Follow-up comparisons indicated that both low and high doses of J60 DREADD ligand increase sucrose consumption in male Gq rats relative to saline control. Similar to chow consumption, we found no effects of J60 ligand in female Gq rats. These results indicate chemogenetic activation of VP GABA neurons increases consumption of both chow and sucrose, selectively in male rats, suggesting that VP GABA neurons impact general consumption in a sex-biased manner, independent of reward type.

### Activation of VP GABA neurons impacts sucrose preference in a sex-dependent manner

Sucrose preference was measured as the percentage of total liquid solution consumed (sum of water and sucrose solutions) that was sucrose solution, during the consumption test. All included rats were noted to have a baseline sucrose preference on habituation days. Gq rats tested for sucrose consumption (n=13, 6 females, 7 males) did not differ in their preference for sucrose across saline (87.51% +/− 1.28), low ligand (87.95% +/− 1.19), or high ligand (88.95% +/− 1.20) test days (Figure 2E). No significant effect of ligand dose (F(2, 22.97) = 0.39, p = .68) or sex (F(1, 10.90) = 1.16, p = .31) was observed in a linear mixed effects model. However, a ligand by sex interaction (F(2, 22.97) = 6.30, p = .007) was observed in the model. Follow up comparisons revealed that low doses of ligand significantly (p = .03) decreased sucrose preference in female rats compared the vehicle controls, whereas both low and high doses of ligand significantly (p = .02, p = .03) increased sucrose preference in male rats. Additionally, sucrose preference following injections of the low dose of J60 was significantly different between male and female rats (p = .02). This effect is aligned with sucrose consumption findings where males consumed more following both doses of ligand and females did not significantly change their sucrose consumption but tended to decrease their consumption compared to vehicle control.

### Activation of VP GABA neurons did not impact water consumption

In Gq rats tested for sucrose consumption and preference, water intake did not differ across saline (1.89 +/− 0.24), low dose (1.94 +/− 0.16), and high dose (2.05 +/− 0.21) test days. No significant effects of injection (F(2, 22) = 0.21, p = .81), or sex (F(1, 11) = 0.71, p = .42), or injection by sex interaction (F(2, 22) = 0.62, p = .54) were found in a linear mixed effects model.

### Inhibition of VP GABA neurons has no significant effect on chow or sucrose consumption

Overall, we observed no significant effects of Gi DREADD manipulations on consumption (Figure 3). In Gi rats tested for chow consumption (n=26, 17 females, 9 males; Figure 3C) we found no significant effects of ligand dose (F(2, 49.29) = 2.15, p = .13) or sex (F(1, 25.17) = 0.01, p = .93), or sex by ligand interaction (F(2, 49.29) = 0.70, p = .50). In Gi rats tested for sucrose consumption (n=15, 8 females, 7 males; Figure 3B and D) we observed a trend toward an effect of J60 ligand in Gi rats, but this effect did not reach significance (F(2,26) = 2.91, p = .07). Additionally, while this effect initially appeared to be driven by a selective decrease in female rats, we observed no effect of sex (F(1,13) = 0.04, p = .85) or sex by injection interaction (F(2, 26) = 0.44, p = .65), and no significant pairwise differences.

**Figure 3.**
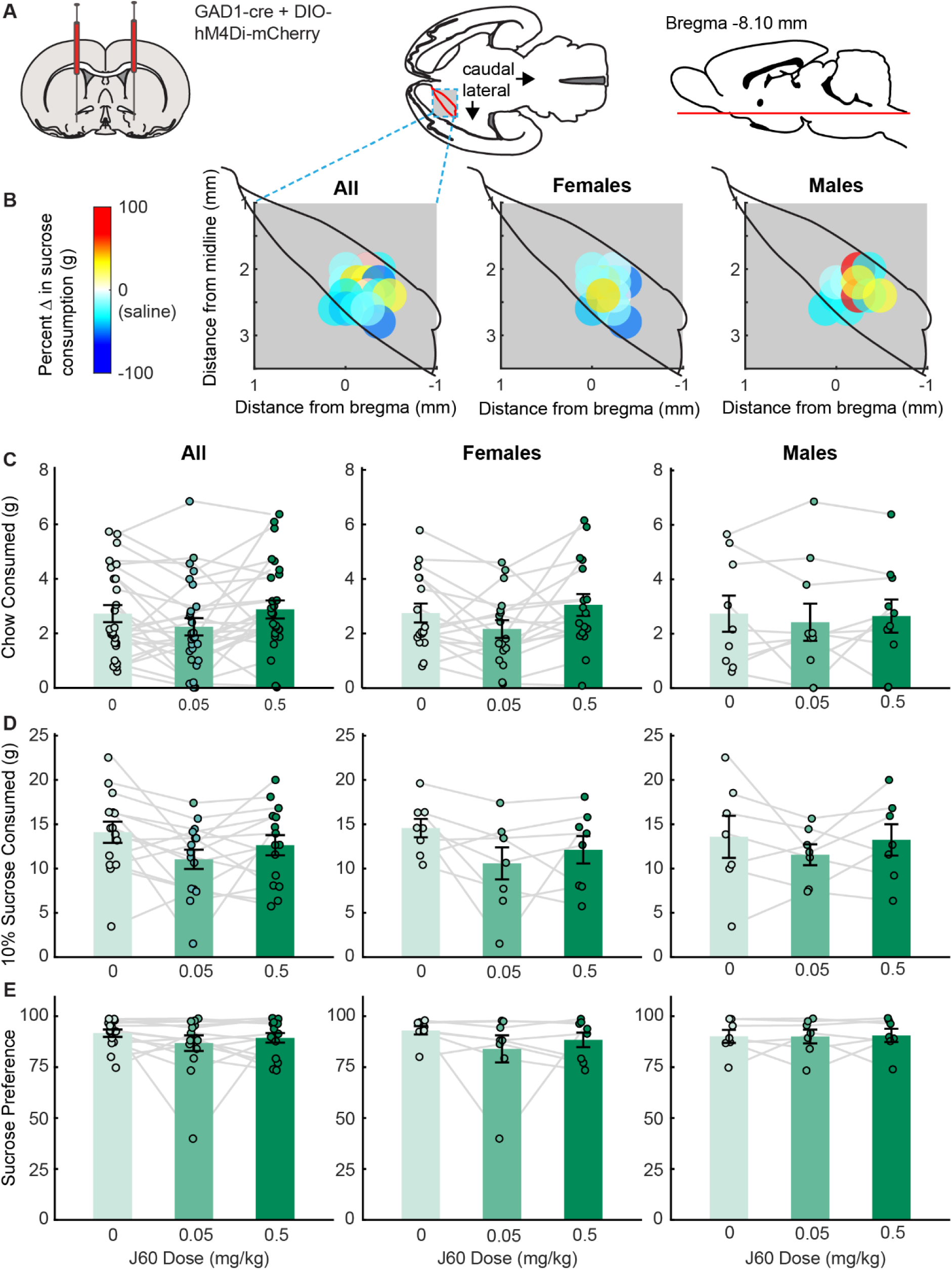
VP GABA inhibition does not significantly affect consumption of chow and sucrose in male or female rats. (A) A mixture of GAD1-cre and DIO-hM4Di-mCherry viruses was injected bilaterally into the VP. (B) Colormaps show viral placement (brightest point of expression) a horizontal cross section of the VP and behavioral changes from baseline following a high dose (0.5 mg/kg) of J60, for all Gi rats (left), female Gi rats (middle), and male Gi rats (right). Chow consumption (C), sucrose consumption (D), and sucrose preference (E) following injections of saline (light green), low dose J60 (medium green) and high dose J60 (dark green) for all Gi rats (left), female Gi rats (middle), and male Gi rats (right). Bar plots and error bars indicate mean +/− SEM. Dots and lines represent individual rats.

### Inhibition of VP GABA neurons did not affect sucrose preference or water consumption

Sucrose preference was measured as described above. All rats were noted to have a baseline sucrose preference on habituation days. Gi sucrose rats (n=15, 8 females, 7 males) did not differ in their preference for sucrose across saline (82.20% +/− 4.37), low ligand (89.13% +/− 2.88), and high ligand (89.46% +/− 2.26) test days. No significant effect of ligand dose (F(2, 60) = 0.69, p = .51) or sex (F(1, 60) = 1.30, p = .26) or ligand by sex interaction (F(2, 60) = 0.63, p = .53) was observed in a linear mixed effect model. Therefore, chemogenetic inhibition of VP GABA neurons did not impact sucrose preference.

### Inhibition of VP GABA neurons did not impact water consumption

Gi sucrose rats (n=15, 8 females, 7 males) did not change their water intake across saline (1.10 +/− 0.23), low dose (1.22 +/− 0.22), and high dose (1.24 +/− 0.24) test days. No significant effects of ligand dose (F(2, 28) = 0.25, p = .78), sex (F(1, 14) = 0.002, p = .96), or ligand by sex interaction (F(2, 28) = 0.29, p = .75) were found in a linear mixed effect model.

### A high dose of J60 DREADD ligand impacts chow consumption but not sucrose consumption in the absence of DREADD receptor

When we examined mCherry control vector rats tested for chow consumption (n=16, 10 females, 6 males; Figure 4C) we found a significant main effect of ligand dose (F(2, 28) = 3.56, p = .04), but no significant effect of sex (F(1, 14) = 1.97, p = .18) or ligand by sex interaction (F(2, 28) = 0.41, p = .67). Follow up comparisons revealed that high dose of ligand significantly (p = .01) increased chow consumption, independent of sex, compared to saline. There was no significant effect of low dose (p = .44) compared to saline, or significant differences between ligand doses (p = .06). This suggests that J60 may act non-selectively at higher doses, consistent with some recent reports (Van Savage and Avegno, 2023). Importantly, when we compared the impact of J60 on chow consumption in male Gq versus control rats, we found a trend towards an interaction between ligand and virus (F(2,26) = 3.22, p = 0.056) suggesting that observed effects in Gq rats are likely due, at least partially, to activation of the DREADD receptor.

**Figure 4.**
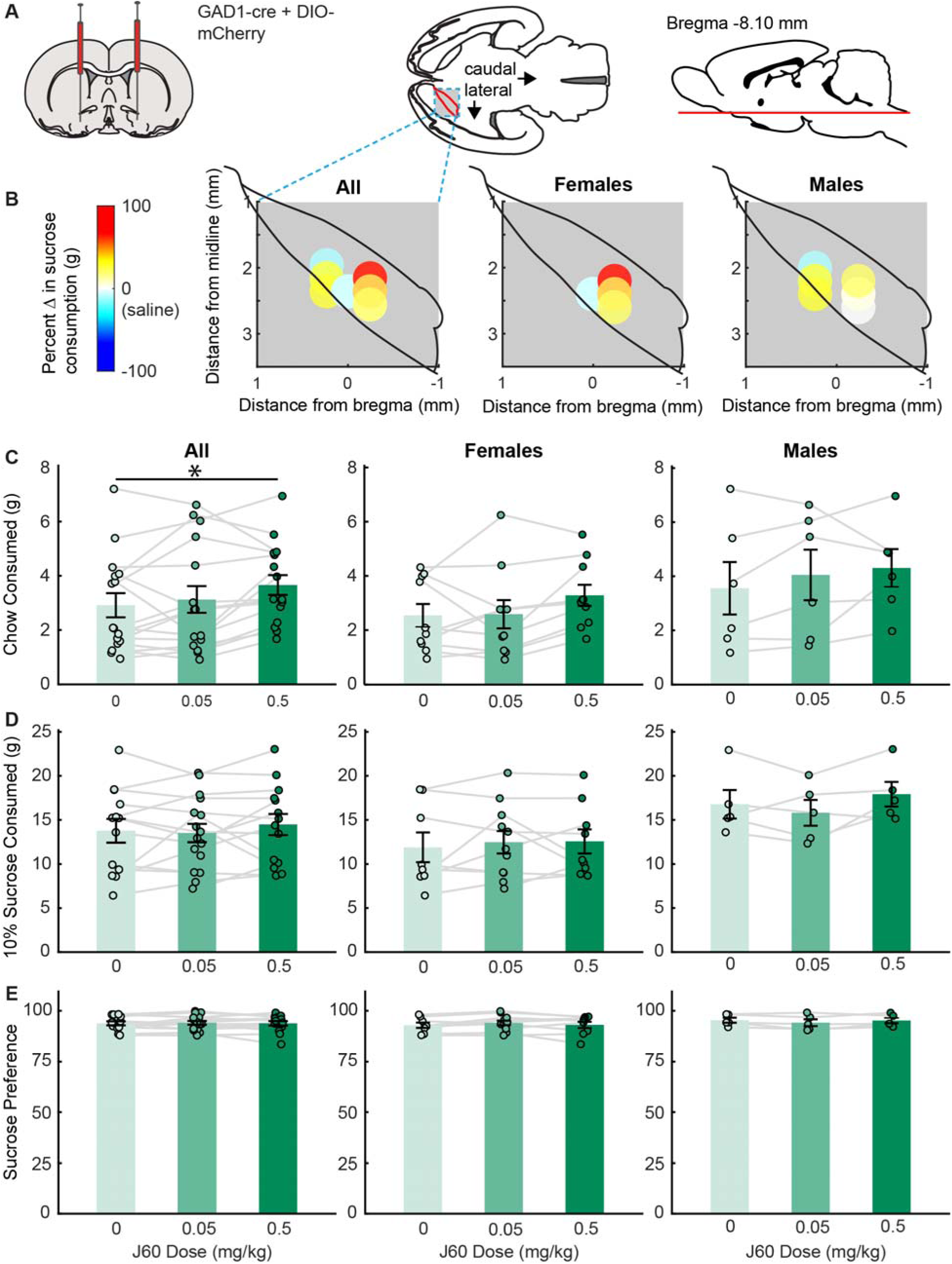
Effect of DREADD ligand in control rats expressing mCherry. (A) A mixture of GAD1-cre and DIO-mCherry was injected bilaterally into the VP. (B) Colormaps show viral placement (brightest point of expression) in a horizontal cross section of the VP and behavioral changes from baseline following a high dose (0.5 mg/kg) of J60 on, for all mCherry rats (left), female mCherry rats (middle), and male mCherry rats (right). Chow consumption (C), sucrose consumption (D), and sucrose preference (E) following injections of saline (light green), low dose J60 (medium green) and high dose J60 (dark green) for all control rats (left), female controls (middle), and male controls (right). Bar plots and error bars indicate mean +/− SEM. Dots and lines represent individual rats. Asterix indicate p-values less than 0.05 for Sidak corrected pairwise comparisons.

When we assessed sucrose consumption (Figure 4B and D) in mCherry control vector rats (n=13, 8 females, 5 males) we found no significant effect of ligand dose (F(2, 25) = 1.18, p = .32), or sex (F(1, 12) = 4.45, p = .06) or ligand by sex interaction (F(2, 25) = 1.78, p = .19). We also found no significant effect of ligand dose (F(2, 25.03) = 0.13, p = .98), sex (F(1, 11.99) = 0.61, p = .45), or ligand by sex interaction (F(2, 25.03) = 1.79, p = .19) for sucrose preference (Figure 4E). When we compared sucrose consumption and preference in Gq and control males we found trends toward an interaction between ligand and virus for both measures (sucrose consumption, F(2,20) = 2.8337, p = 0.082; sucrose preference, F(2,20.97) = 2.73, p = 0.088).

## DISCUSSION

Here we assessed how functional manipulation of the activity of VP GABA neurons impacts consumption of regular chow and 10% sucrose solution in *ad libitum* fed rats. Chemogenetic activation of VP GABA neurons in male rats increased chow and sucrose consumption, as well as sucrose preference. In contrast, chemogenetic activation of VP GABA neurons in female rats failed to increase consumption, and even reduced sucrose preference, suggested sex-biased effects of GABA neuron activation. Inactivation of VP GABA neurons did not significantly affect consumption of either chow, sucrose or water, though we observed a trend toward a decrease in sucrose consumption. Control rats not expressing DREADD receptors increased their chow consumption at the high dose of J60 DREADD ligand, but had no change in sucrose consumption or preference. Importantly, this non-selective effect of the high dose of J60 did not fully account for the effect of J60 in male Gq rats, suggesting that increases in consumption were related to chemogenetic activation, and were specific to male rats.

### Activation of VP neurons impacts chow consumption in a sex-specific manner

When we activated VP GABA neurons in *ad libitum* fed rats we observed significant increases in chow consumption at the high dose of ligand and increases in sucrose consumption and preference at both doses of ligand. This effect differs from findings in Li et al. (2020), where activation of VP GABA neurons had no significant effect on food intake in male mice. Importantly, this prior study tested chow consumption in food-restricted mice, so their null finding may be explained by a ceiling effect on consumption. Alternatively, this difference could be explained by our use of a higher affinity DREADD ligand (Bonaventura et al., 2019).

We were surprised to find sex-biased effects on consumption, though most prior work demonstrating increased consumption following VP disinhibition was conducted in male subjects (Covelo et al., 2014; Stratford and Kelley, 1999; Stratford and Wirtshafter, 2013). Sex-biased effects could be due to the impact of ovarian and other sex-specific hormones on consumption behavior (Alonso-Caraballo and Ferrario, 2019; Butera, 2010; Ma et al., 2020; Xu et al., 2011). Additionally, the internal hunger state of each animal may differ and this could be due to sex hormones affecting metabolism and consumption (Butera, 2010; Xu et al., 2011). While female rats in this study weighed less the male rats at the time of testing, they did not significantly differ in their uncorrected consumption of sucrose under control conditions. Estrogen delivery in ovariectomized female rats causes a significant decrease in food intake when coupled with ghrelin delivery, compared to vehicle and saline controls (Butera, 2010). Estrogen-receptor deletion in most brain regions regulated measures of energy homeostasis, mainly increasing body weight in male and female rats, increasing visceral fat mass and daily food intake in female rats, and decreasing heat production, ambulatory movements and rearing counts in female rats (Xu et al., 2011). These differences suggest that follow-up experiments controlling for effects of circulating sex hormones on metabolism and internal state of the animal would be informative.

### Inhibition of VP GABA neurons has no significant effect on chow or sucrose consumption

When we chemogenetically inhibited VP GABA neurons in *ad libitum* fed rats we saw no significant effect on chow, sucrose, or water consumption, though we saw a trend toward a decrease in sucrose consumption. This is similar to recent findings that sated rats do not significantly change chow-seeking or free access consumption of palatable food (M&Ms) when VP GABA neurons are inhibited (Farrell et al., 2021). In the anteroposterior axis, our virus expression was typically centered on mid to mid-caudal VP locations. It is possible that more rostral or more caudal expression of DREADDs would yield different results, for both Gi and Gq manipulations, based on the respective presence of “coldspots” and “hotspots” in these locations where opioids and orexin agonists have been shown to preferentially impact ‘liking’ and consumption of rewards (Ho and Berridge, 2013; Smith and Berridge, 2007, 2005).

It is likely that there are multiple VP GABA populations (cell type specific and projection specific) that affect consumption, with potentially competing roles in behavior. Simultaneous inhibition of these distinct populations may lead to variable results in consumption tasks. VP GABA neurons project to lateral hypothalmus (LH), a canonical feeding center (Rossi and Stuber, 2018; Stuber and Wise, 2016a), as well as to other areas that are reciprocally connected with LH (nucleus accumbens, lateral habenula, VTA) (Stratford and Wirtshafter, 2013; Root et al., 2015; Stuber and Wise, 2016; Sharpe et al., 2017). Therefore, VP GABA inhibition could be impacting the LH in multiple opposing ways. Chemogenetic “disconnection” of VP GABA neurons and LH, does not significantly impact reacquisition of alcohol seeking in rats, whereas disconnection of VP GABA neurons and neurons in the VTA significantly decreases reacquisition of alcohol seeking (Prasad et al., 2020b). Selective activation or inhibition of VP GABA neurons projecting to these regions may yield dissociable results on food consumption.

The VP GABA neuron population is also composed of heterogeneous cell types such as Npas1, somatostatin, and parvalbumin-containing neurons, among others (Morais-Silva et al., 2023; Prasad et al., 2020a; Pribiag et al., 2021; Zhu et al., 2017). Different VP GABAergic cell type populations can project to distinct areas of the brain involved in reward seeking and consumption behavior (Prasad and McNally, 2016; Zhu et al., 2017; Prasad et al., 2020). Prior research indicates there may be different effects of basal forebrain GABA neuronal subtypes on consumption of highly palatable rewards (Zhu et al., 2017). Additionally, VP GABA neurons have a heterogeneous response to reward paired cues and reward (Stephenson-Jones et al., 2020). Finally, VP GABA parvalbumin neurons have a distinct role acquisition of alcohol seeking behavior (Prasad et al., 2020). Therefore, probing the VP GABA cell types that are involved in food consumption may lead to greater understanding of the neural circuit mechanisms of consummatory behavior.

### Caveats and Future Directions

Here, we used DREADDs to bidirectionally manipulate VP GABA neurons during consumption to examine how activation or inhibition of VP GABA neurons impacts feeding behavior in *ad libitum* fed male and female rats. We found the activation increased consumption selectively in male rats, and that inhibition did not significantly impact consumption. While we observed sex-biased effects, we did not assess estrus cycle, and therefore could not assess how estrus phase impacted our results. Future work examining the impact of estrus cycle, or the removal of circulating hormones, could help to better understand this sex difference. We also did not test how different hunger states affect consumption in our paradigm. Given prior work showing dramatic effects of lesions and non-selective inhibition of VP inhibition on consummatory behavior (Cromwell and Berridge, 1993; Smith and Berridge, 2007), we were surprised to find that inhibition of VP GABA neurons did not significantly impact consumption. It is possible our experiments were underpowered to detect a change with a relatively low effect size, or to detect a sex-biased effect, or that we did not target the regions of VP most responsible for these effects on feeding. Finally, GABA neurons make up a substantial proportion of VP neurons, and more selective manipulation of VP GABA neurons subpopulations based on projection target or more refined cell types will likely yield additional insights into the mechanisms by which VP influences food consumption.

## Acknowledgements

This work was supported in part by National Institutes of Health grant R01DA053208 to JMR, a MnDRIVE Graduate Fellowship in Neuromodulation to AS, and NRSA fellowship F31AA031597 to KEK. Some viral vectors used in this study were generated by the University of Minnesota Viral Innovation Core, and some fluorescence images were taken at the University of Minnesota University Imaging Centers.

